# Constitutive high expression of NOXA sensitizes human embryonic stem cells for rapid cell death

**DOI:** 10.1101/2021.01.13.426566

**Authors:** Richa Basundra, Sahil Kapoor, Emilie Hollville, Nazanin Kiapour, Adriana Beltran Lopez, Nicole Marie Melchiorre, Mohanish Deshmukh

**Author notes:** Correspondence should be addressed to:* **Dr. Mohanish Deshmukh**, Professor, Cell Biology and Physiology, Co-Director, UNC MD/PhD Program, 5102 Mary Ellen Jones Building, 116 Manning Drive, University of North Carolina, Chapel Hill, NC 27599.

## Abstract

Human embryonic stem (hES) cells are highly sensitive to apoptotic stimuli such as DNA damage, which allows for rapid elimination of mutated cells during development. However, the mechanisms that maintain hES cells in the primed apoptotic state are not completely known. Key activators of apoptosis, the BH3-only proteins, are present at low levels in most cell types. In contrast, hES cells have constitutive high levels of the BH3-only protein, NOXA. We examined the importance of NOXA for enabling apoptosis in hES cells. hES cells deleted for *NOXA* showed remarkable protection against multiple apoptotic stimuli. NOXA was constitutively localized to the mitochondria, where it interacted with MCL1. Strikingly, inhibition of MCL1 in *NOXA* knockout cells was sufficient to sensitize these cells to DNA damage, and subsequently, cell death. Our study demonstrates, an essential function of constitutive high levels of NOXA in hES cells is to effectively antagonize MCL1 to permit rapid apoptosis.

**Significance statement:** Human embryonic stem (hES) cells give rise to the entire organism, hence understanding how these cells regulate their survival and death is important. These cells undergo rapid death in response to DNA damage thereby removing mutated cells from the developing embryo. We focused on identifying the mechanism underlying the sensitivity of these cells to DNA damage. We discovered that the protein NOXA is essential for cell death in hES cells. Further, the crucial function of NOXA is to neutralize high levels of antiapototic protein, MCL1, thus enabling hES cells to respond rapidly to DNA damage.

## Introduction

Cells with varied physiological functions exhibit marked differences in their sensitivity to cell death. However, exactly how these differences in apoptotic thresholds are set in cells remain largely unknown. For example, human embryonic stem (hES) cells are highly sensitive to various genotoxic agents such as etoposide, a topoisomerase II inhibitor, UV exposure and γ-irradiation [1-6]. As these cells give rise to the entire organism, their ability to undergo rapid apoptosis following DNA damage is crucial to avoid the propagation of DNA mutations during development. hES cells are also widely used to generate different types of cells for modeling human diseases. Therefore, understanding the mechanisms by which the survival and death of hES cells are regulated is significant for both development and in disease.

Key initiators of apoptosis in cells are the BH3-only members of the BCL2 family proteins [7]. These proteins are activated (*e*.*g*. transcriptional induction, posttranslational modification) by extracellular or intracellular apoptotic stimuli and initiate the cascade of events that result in cell death. Briefly, the BH3-only proteins fall under two categories: - The sensitizer BH3-only proteins (*e*.*g*. NOXA, HRK, BAD) bind to and inactivate the anti-apoptotic BCL2 family proteins such as BCL2, BCLXL, and MCL1. The activator BH3-only proteins (*e*.*g*. BIM, BID, PUMA) bind to and directly activate the pro-apoptotic BCL2 family proteins, BAX and BAK [8]. Thus, activation of BH3-only proteins results in the activation of BAX and BAK which, by forming pores in the mitochondrial outer membrane, induce the release of cytochrome *c* (cyt *c*) into the cytosol to trigger caspases to execute apoptotic cell death.

Interestingly, as the BH3-only proteins consist of many members, they are thought to be functionally redundant. Multiple BH3-only proteins can be engaged in response to a given apoptotic stimuli and several members can act as sensitizer or direct activators to activate BAX and BAK [9-11]. Consistent with the observation that the BH3-only proteins are functionally redundant, the knockout of any single BH3-only protein in mice does not result in complete inhibition of apoptosis. Indeed, with rare exception in specific contexts, deletion of multiple BH3-only proteins (*e*.*g*. triple deletion of *BIM, BID*, and *PUMA*) is necessary to robustly inhibit apoptosis [12].

We previously showed that hES cells undergo apoptosis with DNA damage much more rapidly in comparison to other cell types. For example, while DNA damage typically induces cell death in most cells by 24 to 48 hours, virtually all hES cells die within 5 hours [1]. Multiple factors contribute to this apoptotic priming of hES cells, including increased stabilization of p53 after DNA damage [6], the maintenance of active BAX at the Golgi [1], and the balance between pro-and anti-apoptotic BCL2 family proteins [6]. hES cells are known to express high levels of multiple BH3-only proteins with NOXA being expressed at a strikingly 50-fold higher levels as compared to non-hES cells [13].

This constitutive high expression of NOXA suggests that it could play a crucial role in hES cells. In this study, we examined the function of NOXA in hES cells by generating hES cells that were CRISPR-deleted for *NOXA*. Our results identify NOXA as an essential mediator of DNA damage response and emphasize the importance of NOXA/MCL1 axis in hES cells.

## Methods and Materials

### Cell Culture

Human embryonic stem cell lines H9 (WA09), H14 (WA14) and WA22 were obtained from WiCell Research Institute. hES cells were maintained on hES cell qualified matrigel in mTeSR1 medium. Human normal skin fibroblasts CCD-1079Sk were acquired from ATCC and maintained in DMEM with 10% Fetal Bovine Serum and MEM non-essential amino acids. Cells were maintained at 37°C in 5% CO2.

For cell death assays, cells were seeded at a density of 600,000 cells per well in 6-well plates. Approximately after 24 hours, cells were treated with etoposide or tunicamycin. When MCL1 inhibitors (S63845 and AZD5991) were used, cells were treated with inhibitors in presence or absence of etoposide. Cells were stained with Nuclear Blue to label cell nuclei. Images were captured using Leica DMi8 microscope. Cell death was quantified on the basis of cellular and nuclear morphology, average of more than 800 cells were analyzed per condition. For siRNA transfections, cells were seeded at 350,000 cells per well in 6-well. Approximately after 24 hours, control siRNA (MISSION siRNA Fluorescent Universal Negative Control #1, Cyanine 3) and MCL1 siRNA (GGACUUUUAUACCUGUUAUtt) were transfected using lipofectamine 3000 reagent at 50 nM concentration. Approximately 24 hours post transfection cells, cells were treated with DMSO or etoposide and cell death was quantified as described above.

### Genome editing of *NOXA* locus using CRISPR/Cas9

Two sgRNAs; TCGAGTGTGCTACTCAACTC and TTCTTGCGCGCCTTCTTCCC (Thermo Fisher Scientific) were delivered into H9 cells along with recombinant True Cut Cas9 v2 using the Neon Transfection system (Thermo Fisher Scientific, Cat. No MPK5000). H9 cells (1×10^5^ cells) were electroporated with 33pmols of each sgRNA and 5μg/ml Cas9 diluted in 10 μl buffer R. Cells were electroporated using Neon program 17. Seventy-two hours after electroporation, H9 cells were collected. Editing efficiency was assessed using GeneArt Genomic Cleavage Detection Kit. Cell pools with good indel percentage were dissociated into single cells and seeded on 96-well plates coated with matrigel. After two weeks, the genomic DNA of single-cell colonies was extracted and the *NOXA* gene was amplified using primers flanking the sgRNA target sites. Sanger sequencing confirmed the gene knockout.

### Western blot analysis

For sample preparation, cells were trypsinized using TrypLE select enzyme and floating cells were also collected in experiments where apoptosis was induced. Cell pellets were lysed using RIPA buffer for 30 min on ice followed by sonication (2X, 10s pulses). Samples were centrifuged at 13,000 rpm for 20 min at 4°C. Supernatants were collected and protein quantification was performed using Pierce BCA protein assay kit. Lysates (30-50 μg) were loaded on 12% polyacrylamide gel. After transfer on immobilon FL transfer membrane immunoblotting was performed using specific antibodies (details in supplementary information). Membranes were imaged using LICOR imaging system (Odyssey CLx).

### Immunofluorescence

For immunofluorescence study, cells were seeded on glass coverslips coated with matrigel. For staining with pluripotency markers, cells were incubated at 37°C for approximately 24 hrs and then fixed with 4% paraformaldehyde at 4°C. In drug treatment studies, cells were treated with etoposide for 5 hrs in presence of QVD-OPH (25 mM, SM Biochemicals), thereafter cells were stained for p53, cyt *c*, TOM20. For localization study, cells were first transfected with NOXA-GFP plasmid (Vector Builder) using lipofectamine 3000 reagent as per manufacturers protocol, incubated for 24 hrs and treated with etoposide for 5 hrs in presence of QVD-OPH followed by fixation and staining for MCL1, TOM20 and NOXA-GFP. Hoechst 33342 was used for nuclear staining at all times.

### Cell fractionation

Cells were seeded at ∼ 1.10^6^ cells per well of 6-well plates 24 hrs prior treatment. Cells were incubated in 100 μl of ice-cold CLAMI (Cell Lysis and Mitochondria Intact) buffer (250 mM sucrose, 70 mM KCl, 137 mM NaCl, 4.3 mM Na_2_HPO_4_, 1.4 mM KH_2_PO_4_, 200 μg/ml digitonin, pH 7.2) containing complete protease inhibitor for 5 min on ice. Samples were pelleted at 1,000 g for 5 min at 4°C and supernatants (cytosolic fraction) were collected. Pellets were incubated in 100 μl of ice-cold IP buffer (50 mM Tris-HCl, pH 7.4, 150 mM NaCl, 2 mM EDTA, 2 mM EGTA, 0.2% Triton X100, 0.3% Nonidet P-40) containing complete protease inhibitor for 10 min on ice. Samples were pelleted at 10,000 g for 10 min at 4°C and supernatants (mitochondrial fraction) were collected. Protein concentration were estimated using Pierce BCA protein assay kit and samples containing equal amount of proteins were prepared with SDS-PAGE sample buffer for Western blot analysis.

### Immunoprecipitation

Cells (∼ 5.10^6^ cells) were lysed with 500 µl CHAPS buffer (50 mM Tris-HCl, pH 7.4, 150 mM NaCl, 0.1 mM DTT, 1% CHAPS) containing complete protease inhibitor. Lysates were sonicated and centrifuged at 16,000 g for 5 min at 4°C. Supernatants were precleared with 40 µl protein A/G beads for 30 min at 4°C. Precleared lysates were incubated with 40 µl protein A/G beads preconjugated and crosslinked (5 mM BS3) with 4 µg normal mouse IgG or 4 µg mouse anti-MCL1 IgG_1_ mAb for 4 hrs at 4°C. Beads were washed 3 times with CHAPS lysis buffer before being resuspended with Laemmli buffer for Western blot analysis. **Refer to supplementary material for product details**.

## Results

### hES cells express high level of NOXA

*NOXA* (also called *PMAIP1*) is a BH3-only gene that is known to be induced in response to DNA damage in cells [11, 14, 15]. We examined the basal expression of NOXA in hES cells compared to fibroblasts. Consistent with previous reports, NOXA was present at very high levels in all three hES cell lines (H9, H14 and WA22) as compared to human dermal fibroblasts (Figure 1A) [13, 16]. Since hES cells are known to be primed for apoptosis in response to DNA damage [1] and NOXA is a key mediator of DNA damage-induced apoptosis, we considered the possibility that NOXA could be functionally important for the rapid DNA damage response in hES cells.

**Figure 1.**
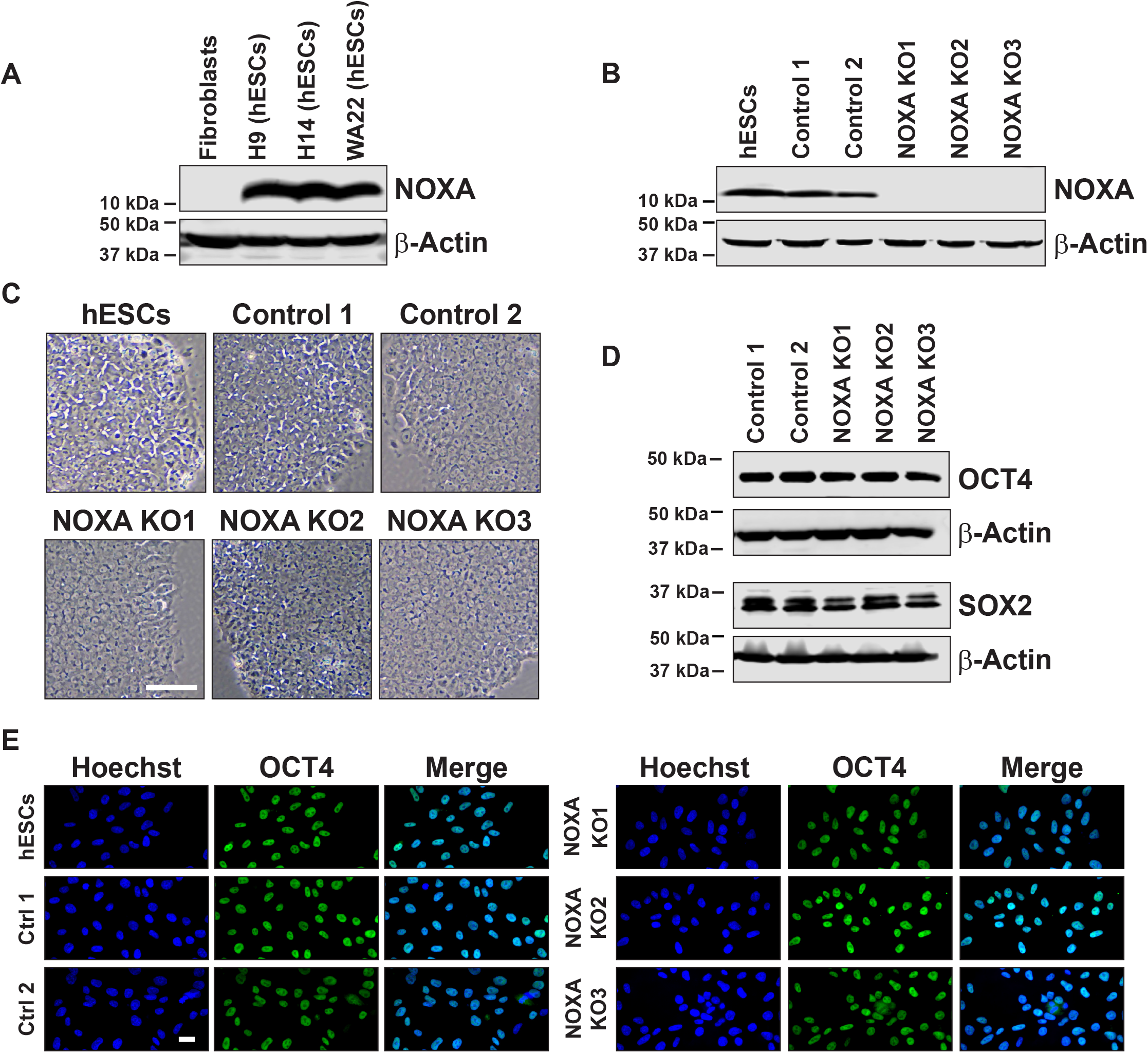
hES cells express high levels of NOXA. (A) Western blot for NOXA expression in human normal skin fibroblasts (CCD-1079Sk), H9 hES cells (WA09), H14 hES cells (WA14) and WA22 hES cells. (B) Western blot for NOXA expression in hES cells (H9), control cells (cells negative for indels) and *NOXA* knockout cells generated by CRISPR/Cas9. (C) Bright field images for hES cells (H9), control and *NOXA* knockout cells. Scale bar 100 μm (representative for all the figures in the panel). (D) Western blot for pluripotency markers OCT4 and SOX2 in control and *NOXA* knockout cells. (E) Immunofluorescence staining for the pluripotency marker OCT4 in hES cells (H9), control and *NOXA* knockout cells. Scale bar 20 μm (representative for all the figures in the panel).

To examine the function of NOXA in hES cells, we utilized CRISPR-Cas9 technology to generate *NOXA* knockout (*NOXA* KO) hES cells. Single cell clone screening was done by sequencing and Western blot to identify hES cells deleted for *NOXA*. Two independent control hES cells and three *NOXA* knockout hES cells were selected for further analysis (Figure 1B).

*NOXA* knockout hES cells maintained their stem cell characteristics, which is evident from their morphology (Figure 1C) and expression of pluripotency markers (*e*.*g*. OCT4, SOX2) (Figures 1D, 1E, S1A and S1B). Thus, *NOXA* deletion alone did not alter the pluripotent status of hES cells.

### *NOXA*-deleted hES cells are resistant to cell death

To examine the functional importance of NOXA in mediating the DNA damage-induced apoptosis in hES cells, we exposed control and *NOXA* knockout hES cells to the DNA damage drug, etoposide. hES cells are very sensitive to DNA damage with even low dose of 1 μM etoposide resulting in the death of 75% of cells within 5 hours of treatment (Figure 2A, B). Strikingly, *NOXA* knockout cells were nearly completely protected under these conditions. Additionally, even at a much higher dose of etoposide (20 μM), *NOXA* knockout cells were markedly protected with greater than 70% cell survival, whereas less than 15% of control hES cells were alive at this concentration (Figure 2A, B).

**Figure 2.**
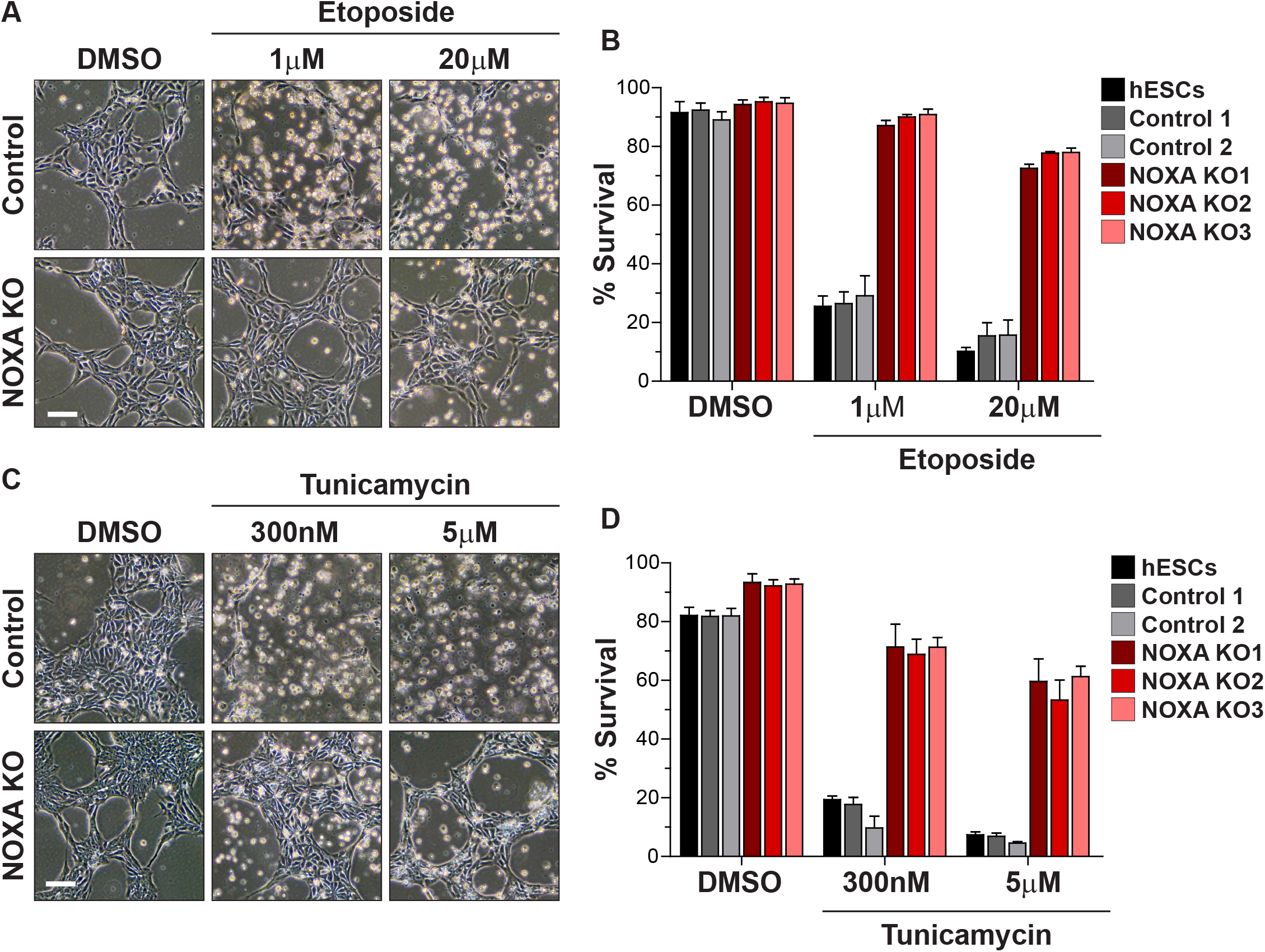
*NOXA*-deleted hES cells are resistant to cell death. (A) Representative bright field images of control and *NOXA* knockout hES cells treated with DMSO or etoposide (1 μM and 20 μM) for 5 hrs. Scale bar 100 μm (representative for all the figures in the panel). (B) Quantification of the survival of hES, control and *NOXA* knockout cells treated with DMSO or etoposide (1 μM and 20 μM) for 5 hrs (n=3 independent replicates; means ± SEM). (C) Representative bright field images of control and *NOXA* knockout hES cells treated with DMSO or tunicamycin (300 nM and 5 μM) for 20 hrs. Scale bar 100 μm (representative for all the figures in the panel). (D) Quantification of the survival of hES, control and *NOXA* knockout cells treated with DMSO or tunicamycin (300 nM and 5 μM) for 20 hrs (n=3 independent replicates; means ± SEM).

To determine if the importance of NOXA in hES cells was specific to DNA damage mediated apoptosis, we examined whether *NOXA* knockout hES cells were also protected in response to endoplasmic reticulum (ER) stress. We focused on ER stress since NOXA has been linked to ER stress-induced apoptosis in various cell lines [17, 18]. Control and *NOXA* knockout hES cells were treated with the ER stress-inducing drug tunicamycin. We found that *NOXA* deletion conferred significant protection at both low (300 nM) and high (5 μM) doses of tunicamycin in hES cells (Figure 2C, D). Similarly, *NOXA* deletion also protected hES cells treated with the broad-spectrum kinase inhibitor staurosporine (data not shown). Together, these results identify NOXA as an essential regulator of apoptosis in hES cells.

### *NOXA* - deficient hES cells induce p53 but do not release cyt *c* after DNA damage

To confirm that NOXA deletion protected against mitochondrial permeabilization in response to apoptotic stimuli in hES cells, we specifically probed for three events that are known to occur in response to DNA damage: i) p53 induction; ii) cyt *c* release from mitochondria; iii) caspase-3 activation. Control and *NOXA* knockout hES cells were treated with 20 μM etoposide for 5 hours and probed for these events. We found that p53 expression was equally induced in both control and *NOXA* knockout cells (Figures 3A and 3B), suggesting that the upstream pathway is unaffected in the *NOXA* knockout cells. However, while cyt *c* was released in approximately 90% of control hES cells, while only 8% of *NOXA* knockout cells showed release of cyt *c* (Figures 3C and 3D). Consistent with the cyt *c* results, while etoposide induced robust cleavage of caspase-3 in control hES cells, only minimal caspase-3 cleavage could be observed in *NOXA* knockout hES cells (Figure 3E). These results confirm that the p53 pathway is functional in *NOXA* knockout hES cells but these cells are unable to release cyt *c* and activate caspase-3 activation after DNA damage.

**Figure 3.**
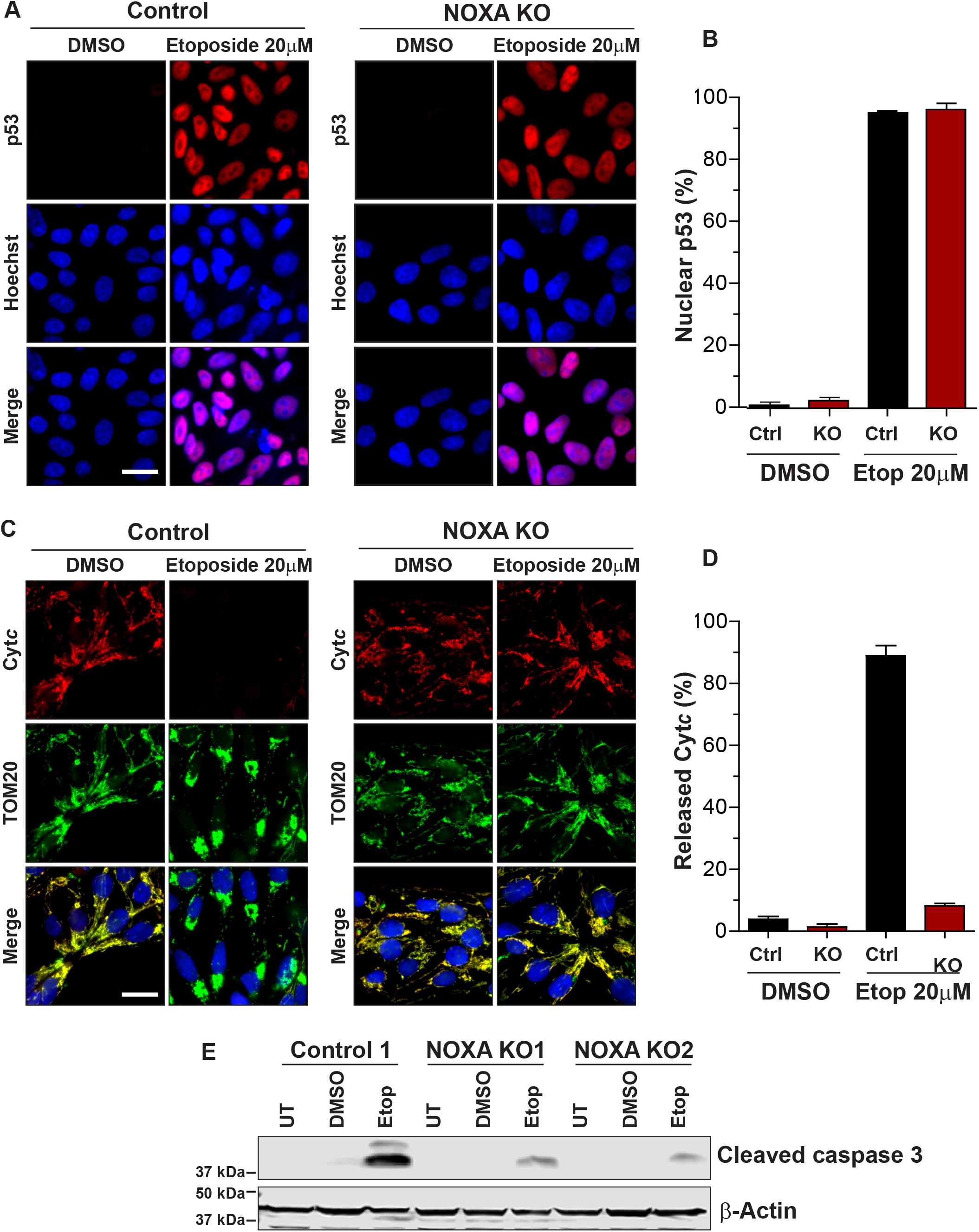
*NOXA*-deficient hES cells induce p53 but do not release cyt *c* after DNA damage. (A) Representative immunofluorescence images of p53 in control and *NOXA* knockout cells treated with DMSO or etoposide (20 μM) for 5 hrs. p53 (red) and Hoechst (blue). Scale bar 20 μm (representative for all the figures in the panel). (B) Quantification of the cells presenting p53 in the nucleus in Figure 3A (n=3 independent replicates; means ± SEM). (C) Representative immunofluorescence images of cyt *c* in control and *NOXA* knockout cells treated with DMSO or etoposide (20 μM) for 5 hrs. Cyt *c* (red), TOM20, mitochondrial marker (green) and Hoechst (blue). Scale bar 20 μm (representative for all the figures in the panel). (D) Quantification of the cells showing cyt *c* released in Figure 3C (n=3 independent replicates; means ± SEM). (E) Western blot for cleaved caspase 3 in control and *NOXA* knockout cells untreated or treated with DMSO and etoposide (20 μM) for 5 hrs (n=3).

### NOXA colocalizes and interacts with MCL1 at the mitochondria in hES cells

Individual BH3-only proteins regulate cell death by either neutralizing the pro-survival proteins (*e*.*g*. BCL2, BCLXL, MCL1) or by directly activating pro-apoptotic proteins (*e*.*g*. BAX, BAK), or both [9]. NOXA is an outlier among the BH3-only proteins because it has limited capacity to interact with multiple BCL2 family proteins and its best defined association is with the pro-survival protein MCL1 [19]. Thus, we examined whether NOXA colocalizes and interacts with MCL1 in hES cells, under normal and apoptotic conditions.

We first tested the localization of NOXA and MCL1 by immunofluorescence. Since we were unable to identify a antibody that specifically detected endogenous NOXA by immunofluorescence in hES cells, we examined the localization of exogenously expressed NOXA-GFP in *NOXA* knockout hES cells. We found NOXA-GFP to be localized to the mitochondria in untreated hES cells as its signal overlapped with the mitochondrial marker TOM20 (Figure 4A). MCL1 staining also overlapped with NOXA and TOM20 staining indicating that both NOXA and MCL1 were present at the mitochondria in untreated hES cells (Figure 4A). Importantly, upon etoposide treatment, the mitochondria appeared aggregated, but both NOXA and MCL1 remained colocalized to the aggregated mitochondria under these conditions (Figure 4A). A similar colocalization of NOXA and MCL1 at the mitochondria was also seen when NOXA-GFP was expressed in control hES cells (Supplementary Figure 2).

**Figure 4.**
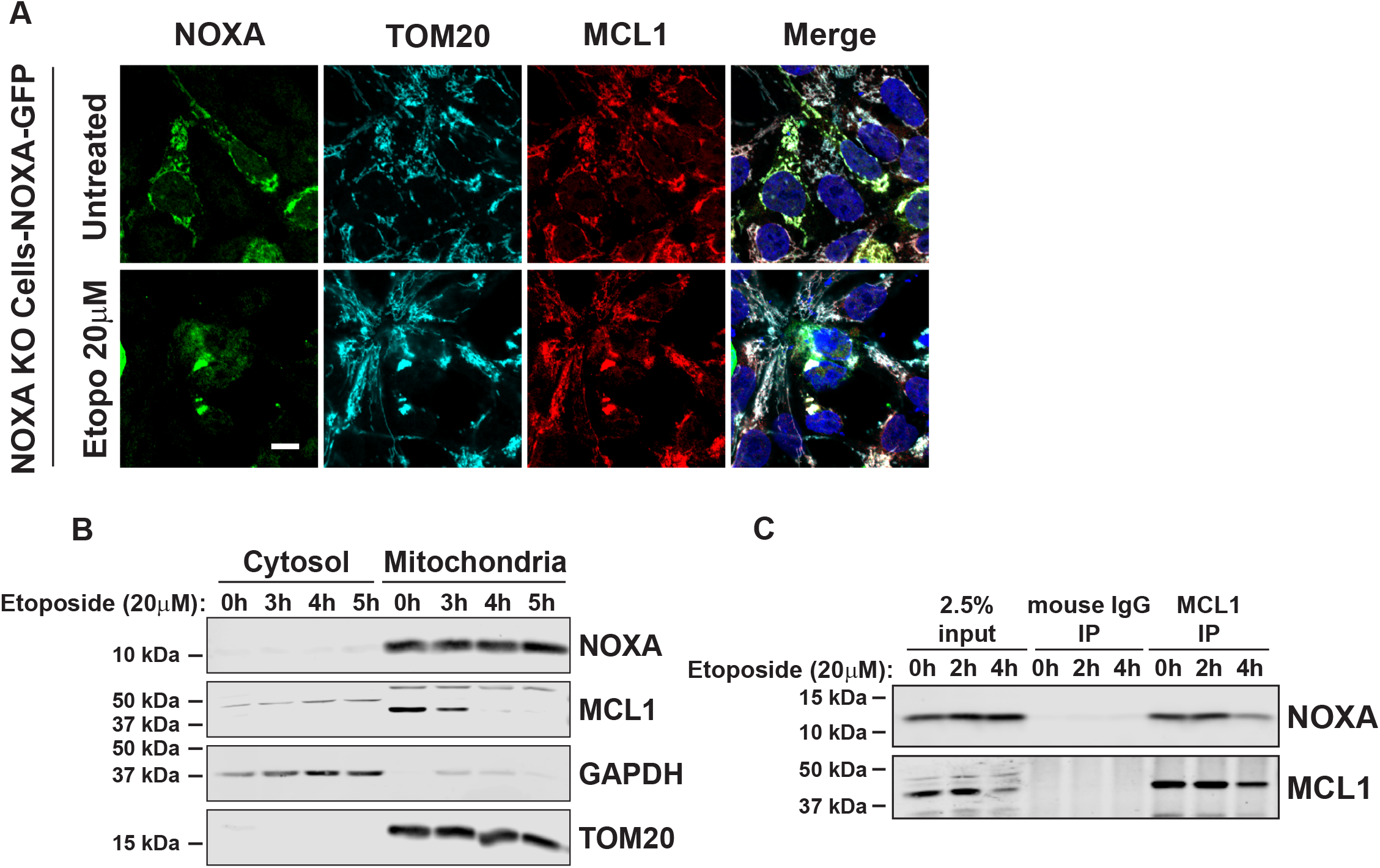
NOXA colocalizes and interacts with MCL1 at the mitochondria in hES cells. (A) Representative images of immunofluorescence staining of NOXA, MCL1, TOM20 in *NOXA* knockout cells transfected with NOXA-GFP plasmid in presence of caspase inhibitor QVD. Cells were either left untreated or treated with etoposide (20 μM) for 5 hrs. Cells were immunostained with antibodies to GFP (NOXA; green), TOM20, (cyan), MCL1 (red) and Hoechst (blue). Scale bar 10 μm (representative for all the figures in the panel). (B) Cytosolic and mitochondrial fractions of cells treated with etoposide (20 μM) for indicated period of time. Fractions were probed for NOXA and MCL1. GAPDH was used as marker for cytosolic fractions and TOM20 as a marker for mitochondrial fraction. (C) Immunoprecipitation of endogenous MCL1 in hES cells treated with etoposide (20 μM) for indicated period of time. Membranes were probed for both NOXA and MCL1.

To confirm the localization of endogenous NOXA and MCL1, we conducted cell fractionation studies in untreated and etoposide-treated hES cells. Analysis of cytosolic and mitochondrial fractions confirmed that both NOXA and MCL1 were present in the mitochondrial fractions of untreated and etoposide-treated hES cells (Figure 4B). Interestingly, MCL1 levels decreased with etoposide treatment, an observation that is consistent with the fact that MCL1 is targeted for degradation in cells undergoing apoptosis [20, 21].

To directly examine whether NOXA and MCL1 interact in hES cells, we conducted immunoprecipitation experiments. Our results show that NOXA is associated with MCL1 in untreated hES cells (Figure 4C). This interaction is initially maintained but subsequently reduced with etoposide treatment, as MCL1 is targeted for degradation (Figure 4B, C). Together these results show that in hES cells, NOXA colocalizes and interacts with MCL1 at the mitochondria under normal conditions, and this interaction is disrupted upon etoposide treatment as MCL1 levels decrease to permit apoptosis.

### MCL1 inhibition permits apoptosis in *NOXA* knockout hES cells

Our results with the *NOXA* knockout cells show that NOXA is essential for apoptosis in hES cells (Figure 2). We considered two possible mechanisms by which NOXA could promote apoptosis in hES cells. While it is generally considered to act as an inhibitor of MCL1, NOXA can also bind to BAX and function as a direct activator of BAX [22]. Thus, the function of NOXA in hES cells could be inhibition of MCL1 or activation of BAX.

To evaluate the specific function of NOXA in hES cells, we examined whether inhibition of MCL1 can restore apoptosis in *NOXA*-deficient hES cells. Control and *NOXA* knockout hES cells were treated with the MCL1 inhibitor S63845 [23] either in absence or presence of etoposide. Our results show that MCL1 inhibition in *NOXA* knockout cells strikingly enabled these cells to undergo apoptosis with DNA damage (Figure 5A, B). To confirm these results, we conducted similar experiments with a second, more recently described, MCL1 inhibitor (AZD5991, [24]). Here too, MCL1 inhibition permitted the *NOXA*-deficient hES cells to undergo apoptosis with DNA damage (Figure 5C). Lastly, as an independent approach to inhibit MCL1, we utilized siRNAs to knockdown MCL1. Our results show that treatment with MCL1 siRNA, but not control siRNA, restored the capability of *NOXA* knockout cells to undergo apoptosis with DNA damage (Figure 5D and S3). Together, these results indicate that the important function of NOXA in hES cells is to inhibit MCL1 and inactivation of MCL1 is necessary for apoptosis to proceed in hES cells.

**Figure 5.**
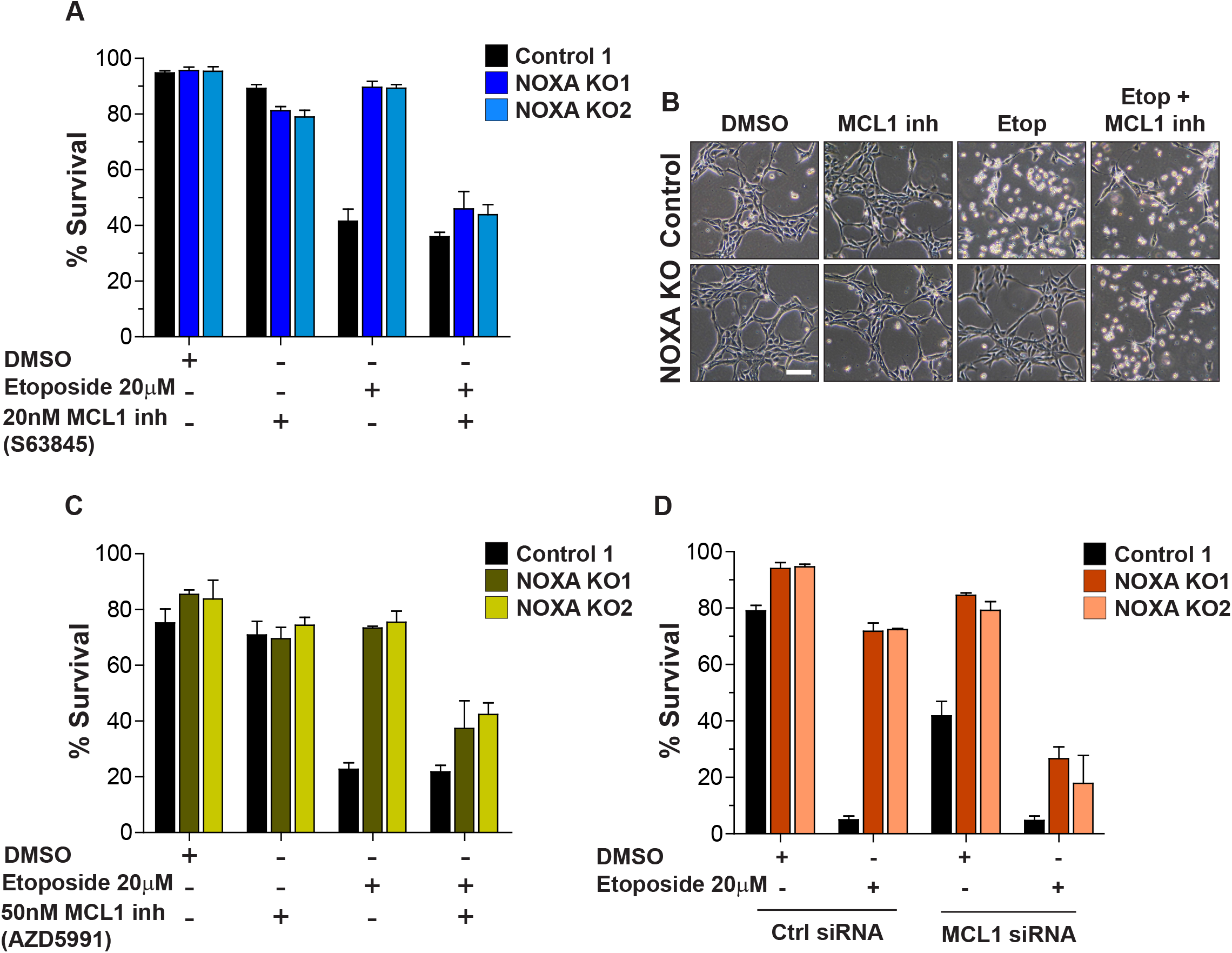
MCL1 inhibition permits apoptosis in *NOXA* knockout hES cells. (A) Quantification of the survival of control and *NOXA* knockout cells in presence of DMSO or MCL1 inhibitor (S63845). Cells were either left untreated or treated with etoposide (20 μM) for 3 hrs, as shown (n=3 independent replicates; means ± SEM). (B) Representative bright field images for Figure 5A. Scale bar 100 μm (representative for all the figures in the panel). (C) Quantification of the survival of control and *NOXA* knockout cells in presence of DMSO or MCL1 inhibitor (AZD5991). Cells were either left untreated or treated with 20 μM etoposide for 5 hrs, as shown (n=3 independent replicates; means ± SEM). (D) Quantification of the survival of control and *NOXA* knockout cells transfected with control siRNA or MCL1 siRNA in presence of DMSO or etoposide (20 μM) for 7 hrs, as shown (n=3 independent replicates; means ± SEM).

## Discussion

In this study, we report that the BH3-only protein NOXA is essential for rapid apoptosis in hES cells. We find that *NOXA* knockout hES cells are resistant to multiple apoptotic stimuli including DNA damage and ER stress. NOXA was also recently identified in a genome-wide CRISPR screen where its deletion was found to improve hES cell survival in response to targeted double stranded breaks [16]. Interestingly, while BH3-only proteins are typically induced with apoptotic stimuli, NOXA is constitutively present at high levels and exists in a complex with MCL1 at the mitochondria even in healthy, untreated, hES cells. Our results show that NOXA-mediated inactivation of MCL1 is a crucial event for apoptosis to occur in hES cells, as MCL1 inhibition permitted apoptosis to occur in *NOXA*-deficient hES cells. Although the BH3-only proteins are generally considered to function redundantly in cells, our results highlight the importance of a single BH3-only protein, NOXA, for apoptosis in hES cells.

*NOXA* is well recognized as a p53-induced gene upon DNA damage in most cells [15]. In hES cells, p53 is induced with DNA damage and required for apoptosis [2]. Strikingly however, its transcriptional activity is not required [3]. This is consistent with the observation that NOXA shows constitutive high expression, and its levels do not substantially increase with DNA damage in hES cells. Interestingly, we have shown previously that p53 is required for the translocation of active BAX from the Golgi to mitochondria [1]. Thus, p53 function in hES cells appears atypical, where it is not required to transcriptionally induce NOXA and activate BAX.

In contrast to other BH3-only proteins that interact with multiple BCL2 family proteins, NOXA primarily binds to only MCL1 with high affinity [14,19]. This ability of NOXA to interact with and inhibit MCL1 is important in contexts where inactivation of MCL1 is necessary for apoptosis to proceed. For example, NOXA is induced in response to multiple apoptotic stimuli including DNA damage, ER stress, hypoxia, and viral infection where it is important for apoptosis [15,25-27]. hES cells not only express high levels of NOXA but also MCL1 [6,28], both of which co-exist at the mitochondria and interact with each other under normal and apoptotic conditions (Figure 4). Importantly, however, MCL1 is targeted for degradation with apoptotic stimuli [21], including in ES cells [27]. Exactly how the apoptotic stimulus promotes MCL1 degradation in hES cells remains unknown. In hematopoietic cells, where NOXA is constitutively expressed [29], or in gastric epithelial cells infected with *Helicobacter pylori*, where NOXA is induced, NOXA is found to be phosphorylated and localized to the cytoplasm [30]. The model proposed is that dephosphorylation of NOXA results in its localization to mitochondria, where it interacts with MCL1 and promotes its degradation. It is unclear if a similar mechanism is engaged in hES cells since NOXA and MCL1 are constitutively at the mitochondria and interact even in untreated conditions. Previous studies have shown that apoptotic stimuli can also cause the phosphorylation of MCL1 that results in its degradation [31]. Thus, we hypothesize that DNA damage likely induces either a posttranslational modification in NOXA or MCL1, or could activate an interacting protein that promotes the degradation of MCL1 in hES cells. Our finding that MCL1 inhibition is sufficient to permit apoptosis in *NOXA*-deficient hES cells indicates that the key function of NOXA in hES cells in the context of cell death is the inhibition of MCL1. These results also indicate that NOXA does not function as a direct activator of the proapoptotic proteins in these cells, since DNA damage-induced apoptosis can proceed in the absence of NOXA with MCL1 inhibition alone.

Examples of cells that constitutively express elevated levels of NOXA are rare. hES cells express NOXA at nearly 50-fold higher levels as compared to other cells [13]. An interesting question here is why are NOXA levels constitutively high in hES cells? In hematopoietic cells where NOXA is also strongly expressed, it functions to increase glucose flux *via* the pentose phosphate pathway [29]. As ES cells are known to be highly glycolytic [32], NOXA could be important for maintaining their high glycolytic state. Additionally, the levels of MCL1 are also high in hES cells where it was found to have non-apoptotic activity for the maintenance of pluripotency. [28]. Even though maintaining high levels of MCL1 is physiologically important for hES cells, the ability of hES cells to undergo rapid apoptosis in response to DNA damage is equally crucial to prevent the propagation of mutations during early embryonic development. Thus, the constitutive high levels of NOXA in hES cells may serve to effectively counteract the anti-apoptotic activity of MCL1 and enable rapid apoptosis. Together, these results highlight the finding that levels and activities of apoptotic proteins are fine tuned to not only to enable their specific non-apoptotic functions in different cell types, but also to set the precise apoptotic thresholds that are physiologically appropriate for those cells.

## Acknowledgements

We thank the Deshmukh lab members for discussions and critical review of this work. We also thank Dr. Anirban Kar, Dr. Ayumi Nakamura, Dr. Vijay Swahari and for reviewing the manuscript. The Neuroscience Microscopy Core Facility, supported, in part, by funding from the NIH-NINDS Neuroscience Center Support Grant P30 NS045892 and the NIH-NICHD Intellectual and Developmental Disabilities Research Center Support Grant U54 HD079124. This work was supported by NIH grant GM118331 to M.D.

## Data sharing statement

Data sharing not applicable to this article as no datasets were generated or analysed during the current study

## Author Contributions

R.B conducted maximum experiments in this study. S.K. helped with the cell death experiments and Western blots. E.H. conducted the cell fractionation, immunoprecipitation experiments and also helped with image aquisition. S.K. and N.K. helped with the quantification of cell death. A.B.L. helped with the generation of the CRISPR knockout hES cells. N.M.M. helped with Western blots. M.D. outlined and supervised the project and M.D. and R.B. wrote the manuscript.

## Supplementary Information

**Figure S1.**
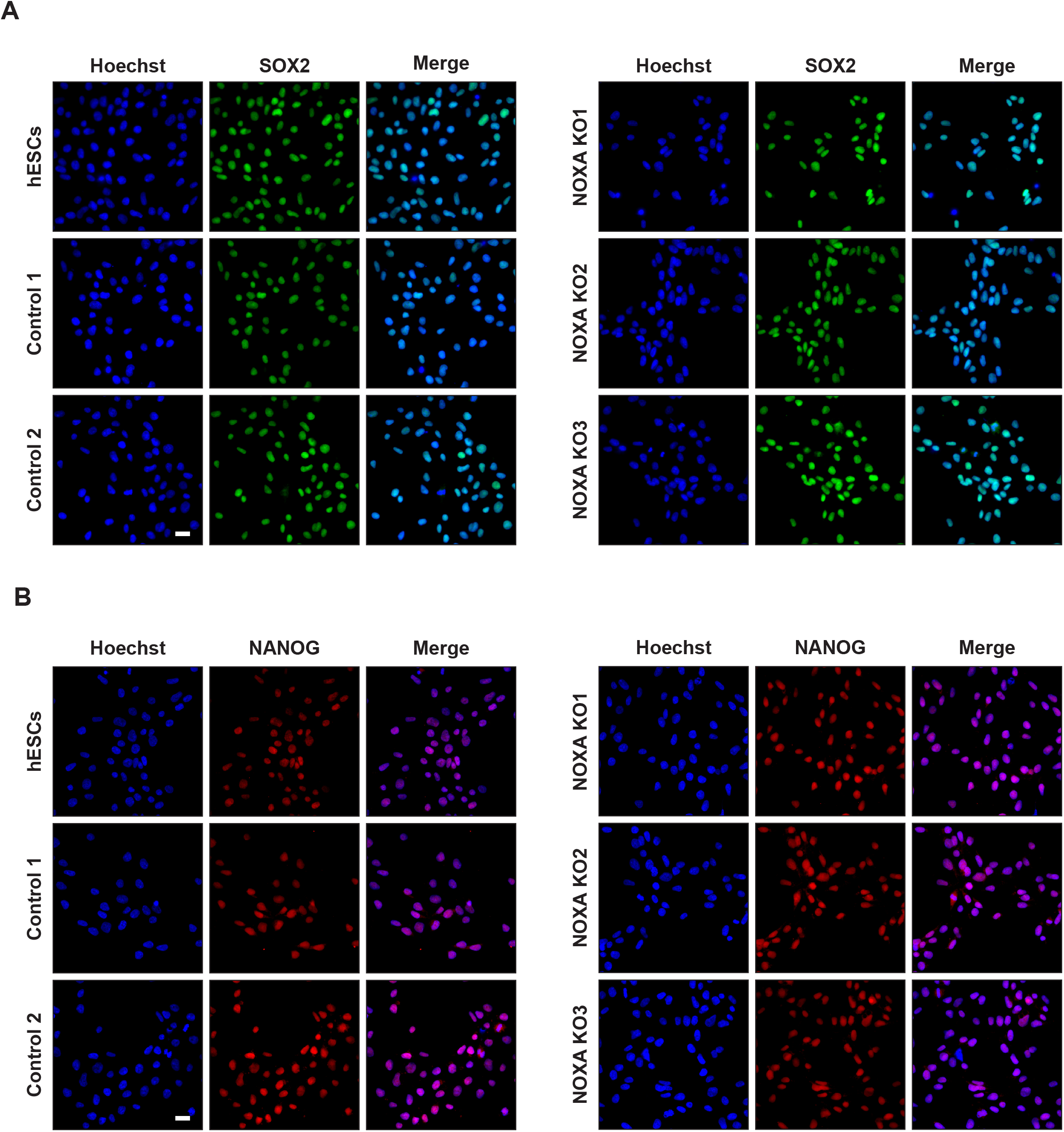
Immunofluorescence staining for pluripotency markers in *NOXA* knockout hES cells. Immunofluorescence staining for pluripotency markers, SOX2 (A) and NANOG (B) in hES cells (H9), control cells (cells negative for indels) and *NOXA* knockout cells generated by CRISPR/Cas9. Scale bar 20 μm (representative for all the figures in the panel).

**Figure S2.**
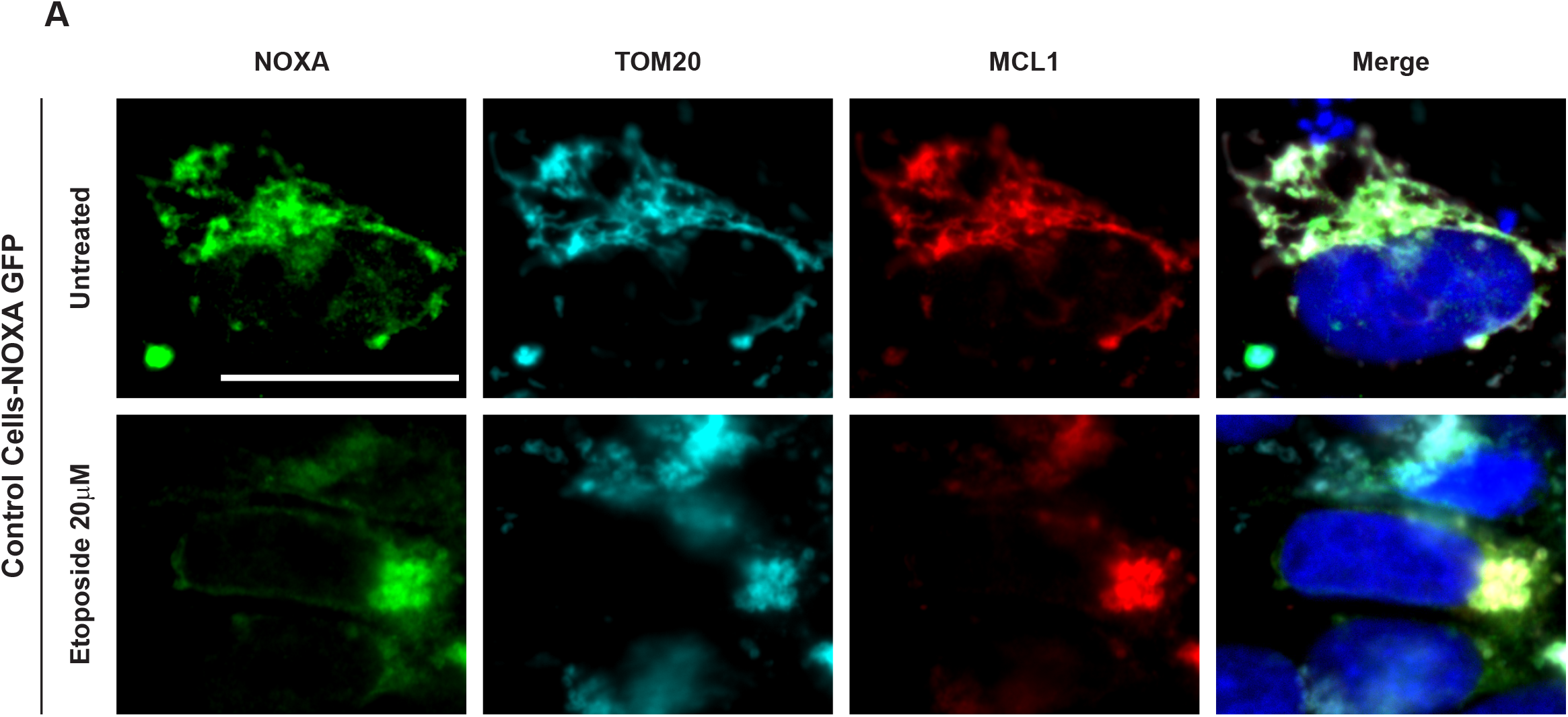
Localization of NOXA and MCL1 in control hES cells. Immunofluorescence staining to show NOXA and MCL1 localization in control cells transfected with NOXA-GFP plasmid, untreated or treated with 20 μM etoposide for 5 hrs. Cells were stained with GFP (NOXA; green), TOM20 (mitochondrial marker; cyan), MCL1 (red) and Hoechst (blue). Scale bar 20 μm (representative for all the figures in the panel).

**Figure S3.**
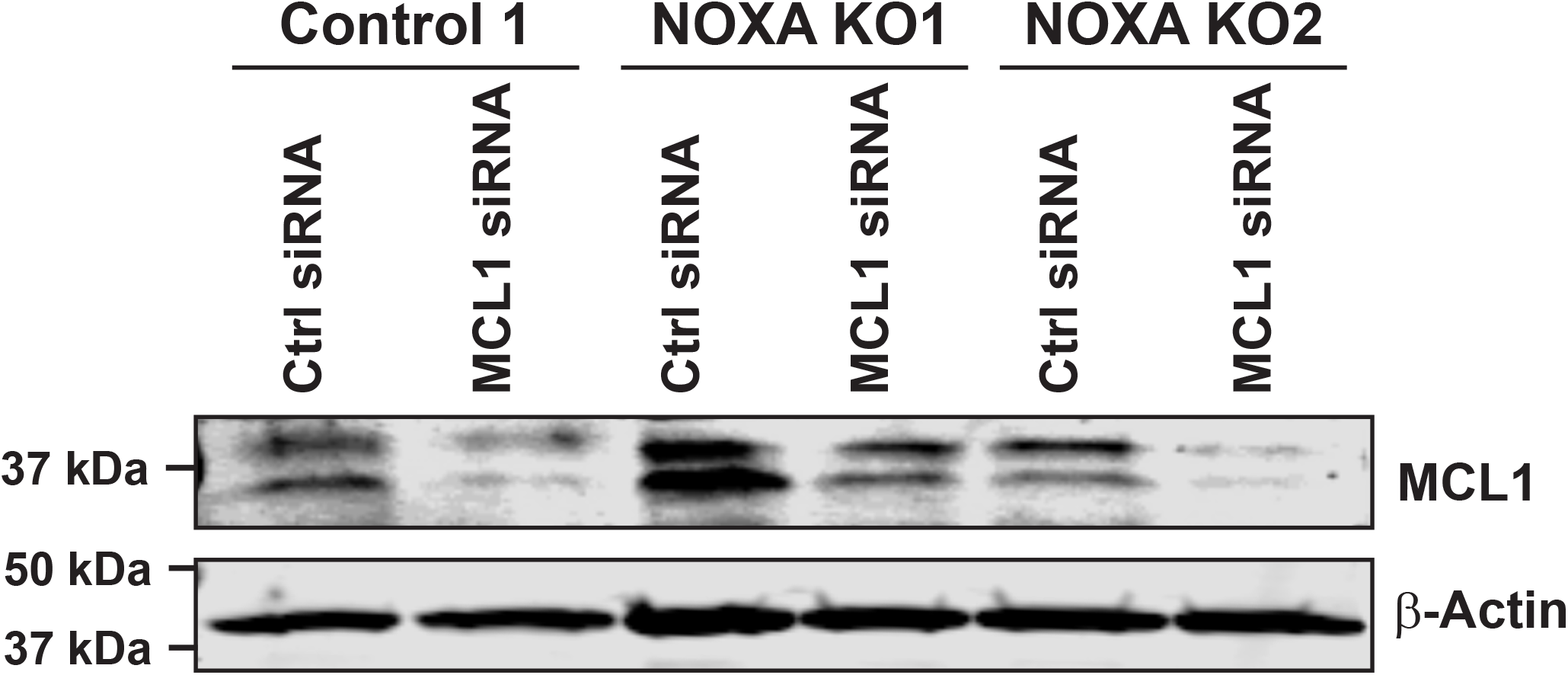
Knockdown of MCL1 in control and *NOXA*-deleted hES cells. Control and *NOXA*-deleted hES cells were transfected with either control siRNA or MCL1 siRNA. Twenty-four hours later, cell extracts were probed for MCL1 and Actin by Western blot analysis.

